# Biomolecular Condensation of Trypsin Prevents Autolysis and Promotes Ca^2+^-Mediated Activation of Esterase Activity

**DOI:** 10.1101/2024.06.01.596986

**Authors:** Chinmaya Kumar Patel, Tushar Kanti Mukherjee

**Affiliations:** Department of Chemistry, Indian Institute of Technology (IIT) Indore, Simrol, Indore 453552, Madhya Pradesh, India

**Keywords:** autolysis, biomolecular condensation, Ca^2+^-binding, esterase activity, macromolecular crowding, phase-separation, trypsin

## Abstract

The presence of Ca^2+^ ions is known to facilitates the biocatalytic activity of trypsin-like serine proteases via structural stabilization against thermal denaturation and autolysis. Herein, we report a new and hidden regulatory role of Ca^2+^ in the catalytic pathways of trypsin and α-chymotrypsin under physiological conditions. We discovered that macromolecular crowding promotes spontaneous homotypic condensation of native trypsin via liquid-liquid phase separation to yield membraneless condensates/droplets in a broad range of concentrations, pH, and temperature. These condensates are stabilized by multivalent hydrophobic interactions between short patches of hydrophobic residues. Importantly, no liquid-to-solid-like phase transition has been observed over a period of 14 days, indicating the structural intrigrity of phase-separated trypsin within the droplets. Structural insights revealed minimal conformational perturbation of trypsin upon phase separation. Interestingly, we found that Ca^2+^ binding in the calcium binding loop reversibly regulates the biomolecular condensation of trypsin and α-chymotrypsin. While Ca^2+^-bound trypsin are ineffective to undergo LLPS to form condensate, its removal facilitates condensation under similar experimental conditions. More importantly, we show that biomolecular condensation effectively prevents autolysis of trypsin at physiological conditions and preserve its native-like esterase activity over a period of 14 days, whereas free trypsin loses 86% of its initial activity. In addition, it has been found that phase-separated trypsin responds to Ca^2+^-dependent activation of its esterase activity even after 14 days of storage while free trypsin failed to do so. Our findings indicate that biomolecular condensates of trypsin and trypsin-like serine proteases act as storage media to prevent autolysis and premature activation, and at the same time preserve their native-like active conformations. The present study highlights an important physiological aspect of biomolecular condensates of trypsin-like serine proteases by which cells can spatio-temporally regulate their biocatalytic efficacy via Ca^2+^-signalling.

## Introduction

Metal ions play key roles in many cellular processes including metabolic regulation,^1^ cell signalling,^2^ energy conversion,^3^ osmotic regulation,^4^ biocatalysis,^5^ and cell membrane potential.^6^ Therefore, living cells precisely regulate the homeostasis of various metal ions for their optimal cellular functions.^7^ Understanding these fundamental regulatory steps for cellular function is important to target various disease associated steps. More fundamentally, metal ions regulate the structural and functional integrity of proteins^8,9^ and enzymes.^10,11^ While the role of metal ions in many metalloenzymes are well documented in the literature, the regulatory role of Ca^2+^ ions on the structural and catalytic activity of serine proteases is poorly understood.

Trypsin is a member of serine proteases with a molecular weight of 23 to 25 kDa. It is involved in the digestive hydrolysis of proteins and exhibits substrate specificity toward positively charged lysine and arginine side chains (Scheme 1A,B).^12–14^ It also exhibits esterase activity in the presence of esters to yield acids and alcohols (Scheme 1B).^13^ Trypsin-like serine proteases find tremendous application in industry and biotechnological fields. This digestive enzyme secreted into the small intestine of animals whereas its proenzyme form, the trypsinogen produced by the pancreas is activated by enterokinase by proteolytic cleavage. Trypsin is produced, stored, and release as the inactive trypsinogen to avoid premature activation which may lead to pancreatic self-digestion and genetic disorders.^15^ Living cells tightly regulate the activity of trypsin to avoid various diseases including pancreatitis, malabsorption, or even cancer.^16^ Trypsin-like serine proteases have a common catalytic triad consisting of histidine-57, aspertate-102, and serine-195 and consist of a double β-barrel fold.^14,17,18^ It contains two calcium (Ca^2+^) binding sites with *K*d values of 0.6 and 16 mM.^19^ While the high affinity (*K*_d_ = 0.6 mM) primary site is located at the calcium-binding loop (CBL) in the N-terminal domain and is necessary for the enzyme’s activity (Scheme 1A),^20^ the low affinity (*K*_d_ = 16 mM) binding site is located in the zymogenic peptide and is necessary for its activation processes via enterokinase. In CBL, Ca^2+^ exhibits octahedral coordination with the side chains of Glu 70 and Glu 80 and the backbone carbonyl oxygens of Asn 72 and Val 75 via electrostatic interactions.^21^ The two other coordination sites are occupied by water molecules. The CBL is formed by residues 69 to 80 in bovine trypsin. Although Ca^2+^ does not play any catalytic role in trypsin activity, the presence of Ca^2+^ is known to facilitates its maximum activity and stability.^22–27^ Earlier, it has been shown that the absence of Ca^2+^ leads to a more flexible structure with considerable increase in the autodigestion of trypsin at alkaline pH. As Ca^2+^ overload or deficiency both leads to various pathological conditions,^15,16^ Ca^2+^ homeostasis is crucial for healthy cells. However, except the stabilizing effect of Ca^2+^ on the activity of trypsin-like serine proteases, no other regulatory role of Ca^2+^ is known for this class of enzymes. Importantly, living cells utilize several regulatory steps to spatio-temporally control the efficacy of various parallel and cascade enzymatic transformations.^28–30^ Understanding these regulatory steps will not only be useful for better catalytic turnover but will help us to design potential therapeutic targets in the pathogenesis of various diseases.

**Scheme 1.**
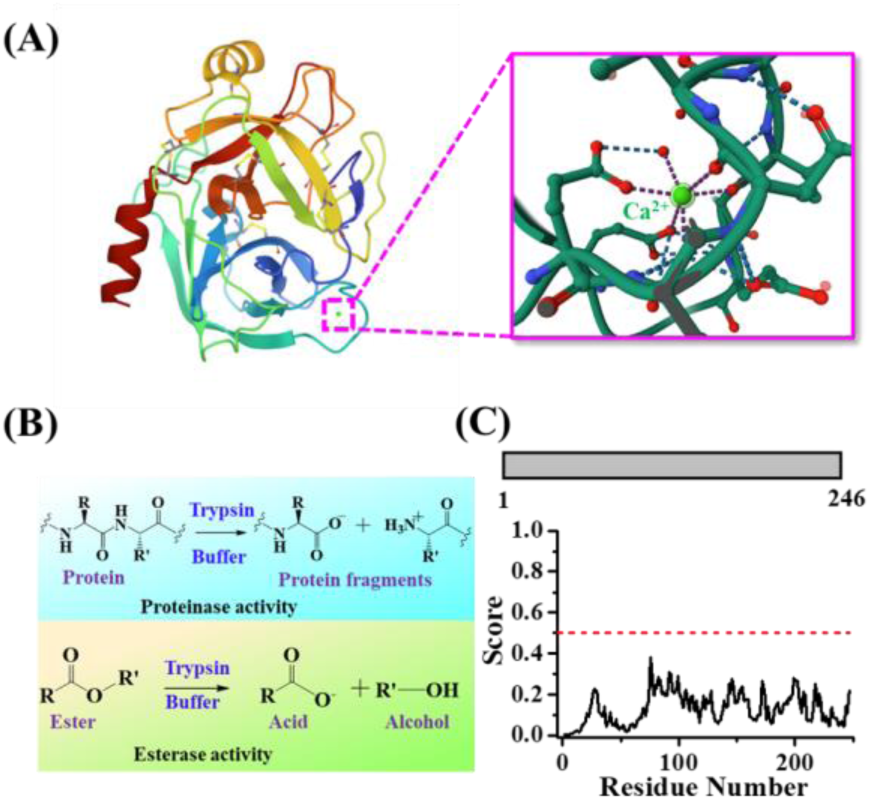
(A) Crystal Structure of Bovine Trypsin (PDB entry 1S0Q). The Enlarged Structure Shows the CBL. (B) Schematics Showing the Proteinase and Esterase Activities of Trypsin. (C) Primary Sequence Analysis of Bovine Trypsin using SMART (upper panel) and IUPred2 (lower panel)

In the present study, we have discovered a unique regulatory role of Ca^2+^ on the activity of bovine pancreatic trypsin and α-chymotrypsin under cell mimicking crowding environment. It is a well-known fact that macromolecular crowding plays crucial role in regulating various fundamental processes including protein folding,^31,32^ DNA replication,^33^ and enzymatic kinetics.^34–44^ Among these, the effect of macromolecular crowding on enzymatic activity is complex and difficult to predict. Lately, it has also been observed that macromolecular crowding promotes liquid-liquid phase separation (LLPS) of many ordered proteins and functional enzymes.^45–49^ In general, proteins containing intrinsically disordered regions (IDRs) with low complexity domains (LCDs) have been shown to undergo LLPS in the absence and presence of inert crowders.^50–64^ LLPS is a spontaneous and thermodynamically favorable liquid-liquid demixing phenomenon for proteins which exhibit weak multivalent attractive intermolecular protein-protein interactions. These interactions involve electrostatic, hydrogen bonding, hydrophobic, dipole–dipole, π–π, and/or cation–π interactions which stabilize the phase-separated condensates/droplets in aqueous medium. Macromolecular crowding promotes these intermolecular interactions via excluded volume effect by lowering the critical concentration of proteins required for phase separation. Due to their highly dynamic and liquid-like nature, biomolecular condensates are highly sensitive towards several factors including animo acid composition, concentration, ionic strength, pH, temperature, pressure, and macromolecular crowding.^45–64^ In the present study, we have investigated the impact of macromolecular crowding on the activity of trypsin in the absence and presence of Ca^2+^ to understand the regulatory role of Ca^2+^ in the esterase activity of trypsin-catalyzed hydrolysis of *p*-nitrophenyl acetate (*p-*NPA) (Scheme 1B). The *p-*NPA hydrolysis by trypsin is a model enzymatic reaction which can be conveniently followed by UV-visible spectroscopy.^34^ Here it is important to mention that not many efforts have been made to understand the effect of crowding on the properties and activity of trypsin at the microscopic level. Earlier, Asaad and Engberts studied the trypsin-catalyzed hydrolysis of *p-*NPA in the absence and presence of PEG and N-tert-butyl acetoacetamide as additives and reported a 2.7-fold decrease in catalytic rate (*k*_cat_) in the presence of PEG.^34^ However, the substrate binding affinity of trypsin (*K*_m_ = 5.57 × 10^-4^ M) was reported to be unaltered in the presence of PEG. It was proposed that the lowering of the enzyme activity is due to an environment effect rather than the crowding induced change in the enzyme conformation. However, no systematic attempts have been made to understand the activity of trypsin in the absence and presence of Ca^2+^ under macromolecular crowding. To better understand the regulatory role of Ca^2+^ on the catalytic activity of trypsin under physiologically relevant heterogeneous crowding environment, we have undertaken the present study.

## Results and Discussion

### Macromolecular Crowding Promotes LLPS of Trypsin-Like Serine Proteases

Recent studies have illustrated that inert synthetic as well as protein crowders can promote LLPS of ordered proteins via promoting multivalent soft protein-protein interactions.^45–49^ Notably, the polypeptide chain of trypsin contains neither any low complexity domains (LCDs) nor any disordered regions as revealed from Simple Modular Architecture Research Tool (SMART)^65^ and IUPred2 sequence prediction algorithms, respectively (Scheme 1C).^66^ Initially, we seek to know whether macromolecular crowding promotes intermolecular protein-protein interactions of trypsin under physiological conditions using confocal laser scanning microscopy (CLSM). All the CLSM measurements were performed inside the aqueous sample chambers fabricated by sandwiching aqueous solution between glass slides and cover slips (Supporting information). Trypsin was fluorescently labeled with fluorescein isothiocyanate (FITC) to visualize under the confocal microscope. To mimic the intracellular crowding, we utilized different inert crowders such as 10% polyethylene glycol (PEG 8000), 12.5% Ficoll 400, 10% dextran 70, and/or 20 mg/mL bovine serum albumin (BSA). Interestingly, the CLSM images of FITC-labeled 5 nM trypsin in the presence of 10% PEG shows the presence of spherical assemblies with uniform green fluorescence (Figure 1A). Notably, no such spherical assemblies are observed in the absence of PEG (Figure S1), indicating the essential role of PEG behind the formation of these assemblies. These assemblies in the presence of PEG display fusion, surface wetting, and dripping phenomena authenticating their liquid-like droplet nature (Figure 1B).^45–47^ Similar liquid-like droplet/condensate formation has been reported previously for a wide range of proteins in the absence or presence of crowders via spontaneous LLPS.^45–64^ Notably, trypsin from porcine and α-chymotrypsin also exhibit LLPS in the presence of 10% PEG to yield liquid-like condensates (Figure S2). To know whether the droplet formation phenomenon of trypsin is specific for PEG or a general crowding effect, we recorded the CLSM images of trypsin in the presence of 12.5% Ficoll 400, 10% dextran 70, and 20 mg/mL BSA individually. Remarkably, irrespective of the nature of the crowders, we observed uniform spherical droplets suggesting a general crowding effect (Figures 1C and S3). The phase diagram of the binary mixture of trypsin and PEG reveals that the critical concentration of trypsin required for phase separation decreases with increase in the PEG concentrations (Figure 1D). This trend can be explained by considering excluded volume effect as high concentration of PEG promotes enhanced protein-protein interactions at lower concentration of trypsin owing to the reduction in the effective volume available to the individual trypsin molecule. Moreover, hydrophilic crowder PEG is known to promotes phase separation of biomolecules via partial dehydration of the polypeptide chain without partitioning into the dense droplet phase.^67^ To know whether crowders are excluded from the droplet phase during LLPS, we performed phase separation assay with unlabeled trypsin in the presence of FITC-labeled PEG (mPEG-NH_2_, MW 5000) and BSA. While the phase contrast images show the presence of well-dispersed spherical droplets, the fluorescence images reveal that the labeled PEG and BSA are completely excluded from the droplet phase as no visible fluorescence signals are detected from inside the individual droplets (Figures 1E,F and S4). These findings authenticate that the LLPS of trypsin in the presence of inert crowders is homotypic in nature similar to other proteins.^45–49,54^ The mean size of trypsin condensates increases progressively upon increase in the concentrations of trypsin at a fixed concentration of PEG (10%) (Figure 1G). This observation can be explained by considering dominant protein-protein interactions as well as extensive number of coalescence/fusion events at high trypsin concentration. Similarly, at a fixed concentration of trypsin (5 nM) and PEG (10%), the mean size of these condensates increases as a function of incubation time at 37 ℃ due to coalescence and Ostwald ripening phenomena (Figure 1H). Taken together, our observations indicate that trypsin and trypsin-like serine proteases undergo spontaneous homotypic phase separation via formation of liquid-like biomolecular condensates under cell mimicking crowding environment through intermolecular protein-protein interactions.

**Figure 1.**
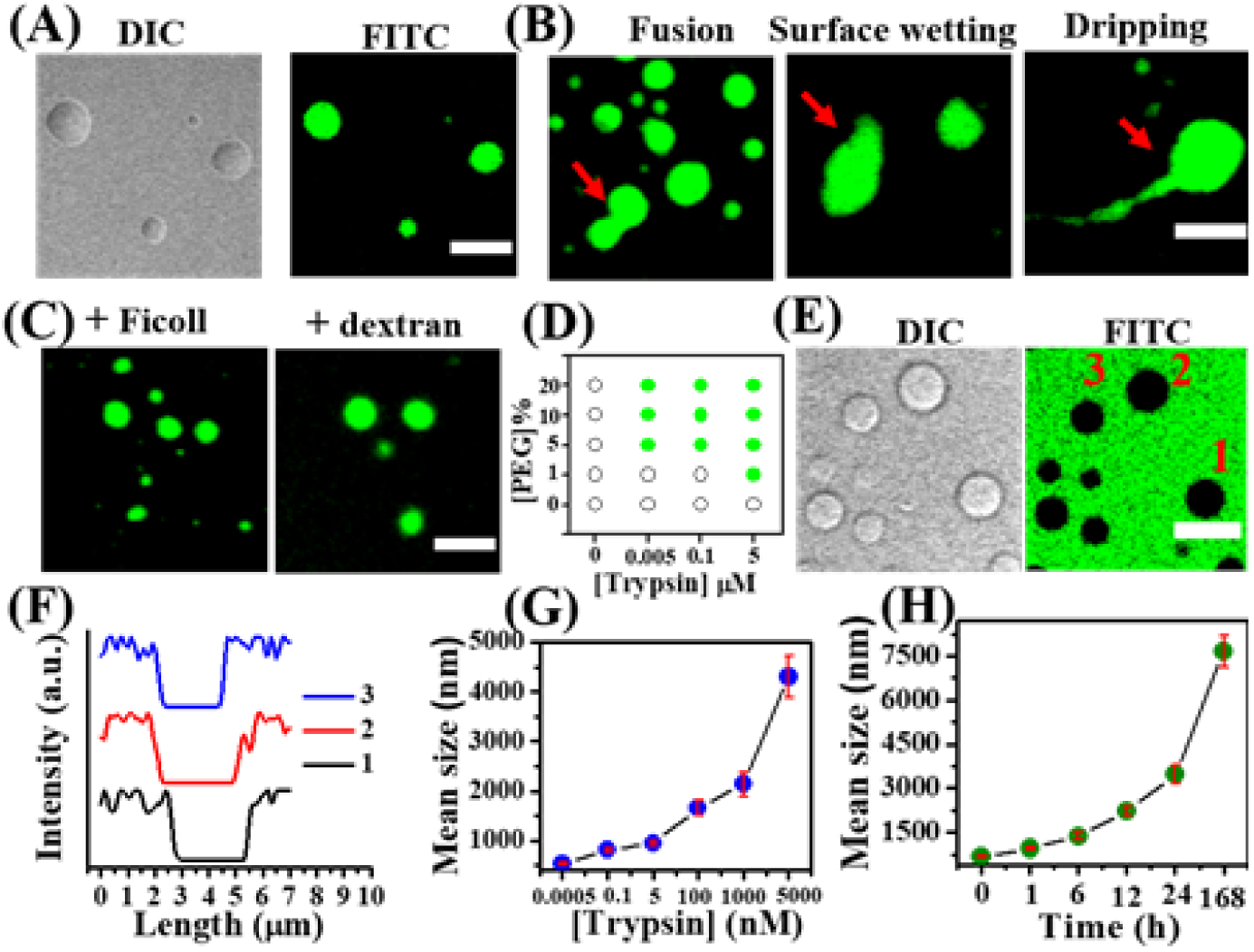
Confocal images of FITC-labeled 5 nM trypsin showing (A) droplet formation in the presence of 10% PEG 8000 upon 1 h incubation in pH 7.4 PBS at 37 °C, (B) fusion, surface wetting, and dripping events (red arrows) of liquid-like droplets, and (C) droplet formation in the presence of 12.5% Ficoll 400, 10% dextran 70. (D) Phase diagram of trypsin as a function of PEG concentration. (E) Confocal images of unlabeled trypsin in the presence of FITC-labeled 10% mPEG-NH2 (MW 5000), and (F) corresponding intensity line profiles of three representative droplets. Variation of mean droplet size of trypsin as a function of (G) trypsin concentrations and (H) incubation time in the presence of 10% PEG 8000 in pH 7.4 PBS at 37 °C. The data points represent the mean ± s.e.m. from three independent measurements. Scale bars correspond to 5 µm.

Next, we seek to address the conformational stability of phase-separated trypsin inside the condensates using circular dichroism (CD) and Fourier-transform infrared spectroscopy (FTIR). Notably, the aqueous dispersion of trypsin in the presence of 10% PEG remains isotropic in nature even at 100 µM protein concentration and no visible turbidity appears within 14 days of incubation at 37 ℃ (Figure 2A). More importantly, no liquid-to-solid-like phase transition has been observed within 14 days of incubation suggesting colloidal and structural integrity of the phase-separated trypsin inside the droplet phase. To know the conformational insight of the phase-separated trypsin in the presence of 10% PEG, we recorded the CD spectra as a function of incubation period. Trypsin in the absence of PEG shows a distinct CD peak at 210 nm with a weak shoulder at 225 nm (Figure 2B).

**Figure 2.**
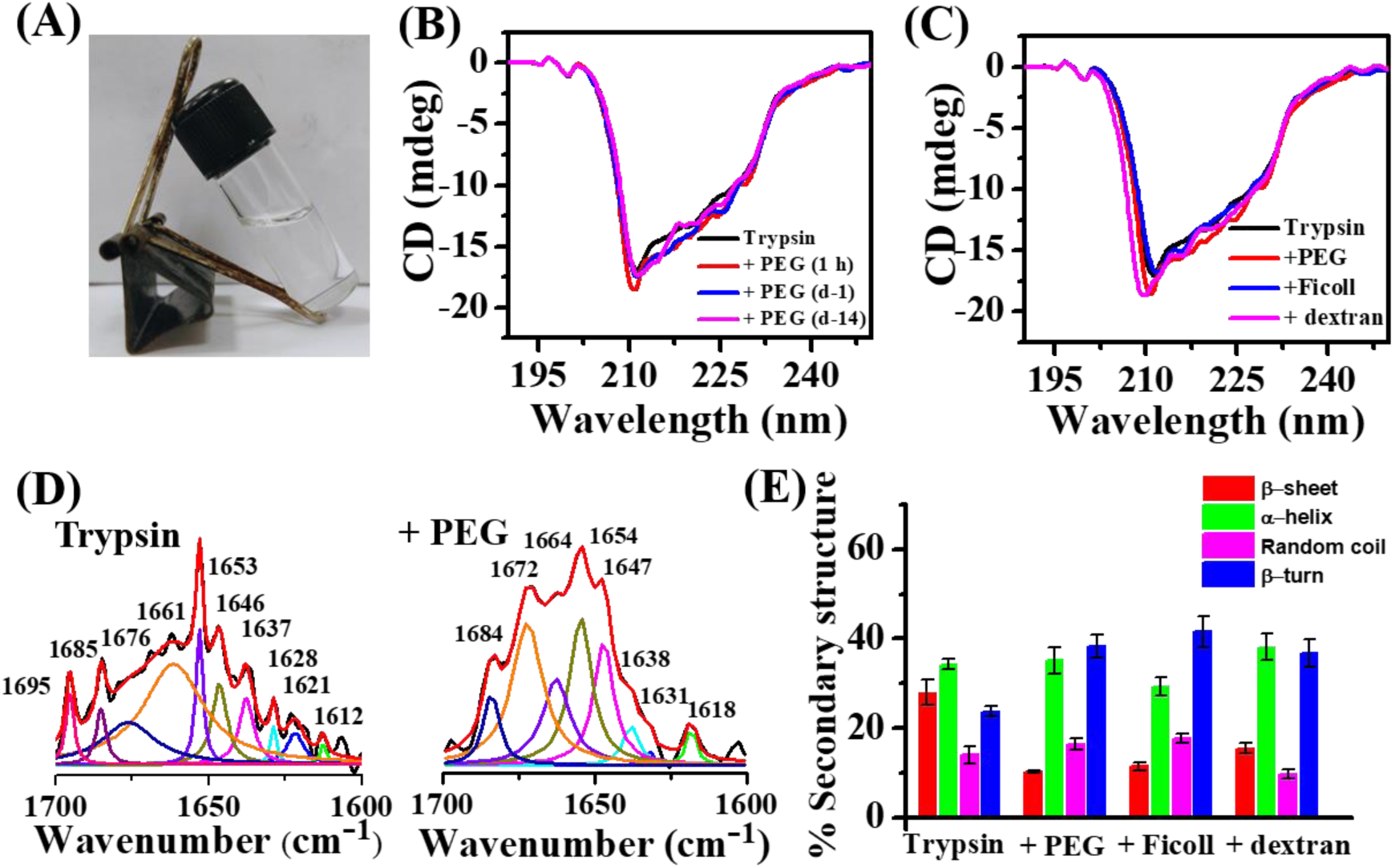
(A) The daylight photograph of aqueous dispersion of 100 µM trypsin in the presence of 10% PEG 8000 upon 14 days of incubation at 37 ℃. CD spectra of 100 µM trypsin in the absence and presence of (B) 10% PEG 8000 and (C) different crowders. (D) Deconvoluted FTIR spectra of 100 µM trypsin in the absence and presence of 10% PEG 8000. (E) Secondary structure contents of trypsin in the absence and presence of different crowders estimated from deconvoluted FTIR spectra. The data points represent the mean ± s.e.m. from three independent measurements. All the solutions were prepared in pH 7.4 PBS and kept at 37 °C.

No appreciable spectral changes have been observed over a period of 14 days in the CD spectrum of trypsin, authenticating the conformational integrity of the phase-separated trypsin (Figure 2B). On the other hand, the native secondary structure of trypsin remains almost unaltered in the presence of different crowders upon phase separation as revealed from CD measurements (Figure 2C). FTIR measurements were utilized to extract the secondary structure contents of trypsin in the absence and presence of different crowders using spectral deconvolution method (Figure 2D,E). The deconvoluted FTIR spectrum of trypsin reveals 28% β-sheet, 34% *α*-helix, 14% random coil, and 24% β-turn in its secondary structure (Figure 2E). A marginal change in the FTIR spectrum of trypsin has been observed in the presence of different crowders (Figures 2D and S5). Spectral analyses reveal that the α-helix content remains unaltered to 30–38%; however, the β-sheet content decreases to 10–16% with concomitant increase in the β-turn content to 37–42% in the presence of different crowders (Figure 2E). These findings indicate that although trypsin does not undergo significant conformational alteration upon phase separation in the presence of different crowders, the marginal local conformational fluctuation of trypsin is quite evident possibly due to intermolecular soft protein-protein interactions. Next, we seek to know the nature of these soft intermolecular interactions.

### Role of pH and Temperature on the LLPS

Proteins often interact with neighbouring proteins through various noncovalent interactions and inert crowders are known to promote these interactions via excluded volume effect.^45–49,54^ These weak but multivalent protein-protein interactions often drive the phase separation of biomolecules and stabilize the phase-separated condensates. To know the nature of the intermolecular interactions in the present system, we first studied the influence of pH and temperature on the phase separation propensity of trypsin. The pH-dependent phase separation assays were performed in the pH range of 3.5–10 (Figure 3A). While trypsin undergoes LLPS in the pH range of 5–10, no phase separation has been observed at pH 3.5. The reported isoelectric point (pI) of bovine pancreatic trypsin is close to 10.5 and hence it remains as a cationic form in the pH range of 3.5–10.^68^ Therefore, the spontaneous phase separation observed at pH 9 and 5 suggests that the long-range electrostatic interactions play negligible role in driving the LLPS of trypsin. The lack of any phase separation at pH 3.5 could be due to the conformational alteration of native trypsin which prevents the intermolecular protein-protein interactions. To authenticate this argument, we recorded the CD and FTIR spectra of trypsin at pH 3.5 and compared with those at pH 10. Both the CD and FTIR data reveal significant alteration in the secondary structure of trypsin at pH 3.5 compared to those at pH 10 (Figure 3B,C). A significant decrease in the α-helix content from 36 to 20% with concomitant increase in the random coil and β-sheet content from 12 to 20% and 26 to 32%, respectively are evident upon decreasing the pH from 10 to 3.5 (Figure 3C). These conformational alterations effectively modulate the favorable intermolecular protein-protein interactions responsible for LLPS of trypsin and ultimately prevent the phase separation. Similar pH-dependent modulation of phase behavior has been reported earlier for various other proteins.^45–48^

**Figure 3.**
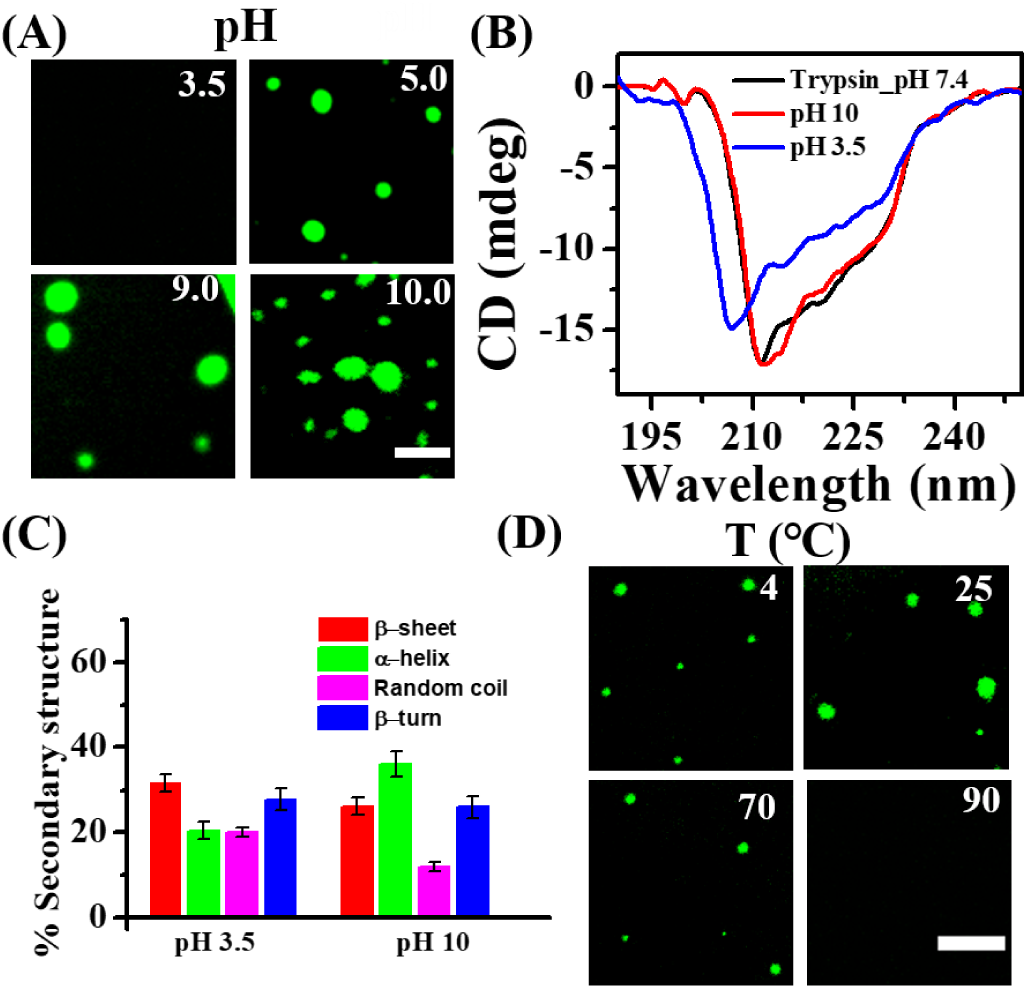
(A) Confocal images of FITC-labeled trypsin (5 nM) droplets at different pH. (B) CD spectra of 100 µM trypsin at different pH. (C) Secondary structure contents of trypsin at pH 3.5 and pH 10 estimated from the deconvoluted FTIR spectra. The data points represent the mean ± s.e.m. for three independent measurements. (D) Confocal images of FITC-labeled 5 nM trypsin in the presence of 10% PEG 8000 at various temperatures. The scale bars correspond to 5 μm. All the samples were incubated at 37 °C in pH 7.4 PBS for 1 h.

Next, to decipher the role of temperature on the phase behavior of trypsin, we varied the solution temperature in the range of 4–90 ℃ and followed the phase separation assay using CLSM (Figure 3D). Temperature in general shows a profound impact on the feasibility of biomolecular condensation as the phenomenon is mainly governed by multivalent weak interactions. Biomolecular condensation can follow either upper critical solution temperature (UCST)^45–47^ or lower critical solution temperature (LCST)^51^ or both.^55^ The phase separation assays were performed by mixing 5 nM trypsin with 10% PEG in pH 7.4 PBS and the binary mixtures were equilibrated at various temperature for 1 h (Figure 3D). The presence of well-dispersed droplets of trypsin is evident in the temperature range of 4–70 ℃; however, condensation is completely inhibited at 90 ℃, suggesting that the LLPS of trypsin follows UCST profile. The present UCST profile of trypsin condensation suggests that the LLPS trypsin is mainly driven by enthalpically favourable (Δ*H* ˂ 0) intermolecular interactions under macromolecular crowding. Various other proteins and functional enzymes have also been shown to undergo LLPS using UCST profile.^45–47^ On the other hand, biomolecular condensations which are feasible only at elevated temperature follow LCST profiles and are mainly driven by entropy.^51^ Here it is important to mention that biomolecular condensation is energetically favorable (Δ*G* ˂ 0) only when the enthalpy gain through multivalent protein-protein interactions overcomes the entropy of mixing according to the Flory–Huggins theory.^69,70^ The present biomolecular condensation of trypsin is only feasible under macromolecular crowding as it promotes intermolecular interactions which help to overcome the unfavorable entropy loss associated with the de-mixing process. Taken together, our findings reveal that the solution pH and temperature both can effectively regulate the biomolecular condensation of trypsin via modulation of various intermolecular protein-protein interactions at the microscopic level.

### Intermolecular Interactions Behind LLPS

Biomolecular condensation is often driven by various intermolecular interactions which includes electrostatic, hydrophobic, π-π, cation-π, and/or hydrogen bonding. To recognize the true nature of intermolecular interactions in the present system, we performed different sets of controlled experiments by varying the concentrations of various additives. First, we varied the salt (NaCl) concentrations from 50 mM to 3 M to know whether electrostatic interactions play any role in the phase separation of trypsin as high salt concentration mask the electrostatic interactions. The salt-dependent phase separation assays were followed under a confocal microscope. Notably, irrespective of the NaCl concentration, trypsin undergoes LLPS to yields uniform spherical droplets (Figure 4A), suggesting electrostatic interactions play no role in the LLPS of trypsin. Further, to know the hydrophobic protein-protein interactions.^45–47^ It is observed that phase separation of trypsin is feasible up to 3% of 1,6-hexanediol; however, it is prevented at 6% of 1,6-hexanediol whether hydrophobic interactions play any role in the present phase separation, we varied the concentrations of 1,6-hexanediol and sodium thiocyanate (NaSCN), which are known to mask (Figure 4B). Similarly, the trypsin condensation is not feasible at and above 1.0 M NaSCN (Figure 4C). Furthermore, addition of a kosmotropic salt, ammonium sulfate (NH_4_)_2_SO_4_ favors the condensation of trypsin, as revealed from the formation of larger-sized droplet in the CLSM images (Figure 4D). On the other hand, addition of adenosine triphosphate (ATP), an electrostatic disruptor has no effect on the phase separation of trypsin in the presence of crowders (Figure 4E). All together, these microscopic observations substantiate the active role of multivalent hydrophobic interactions behind the LLPS of trypsin in the presence of inert crowders.

**Figure 4.**
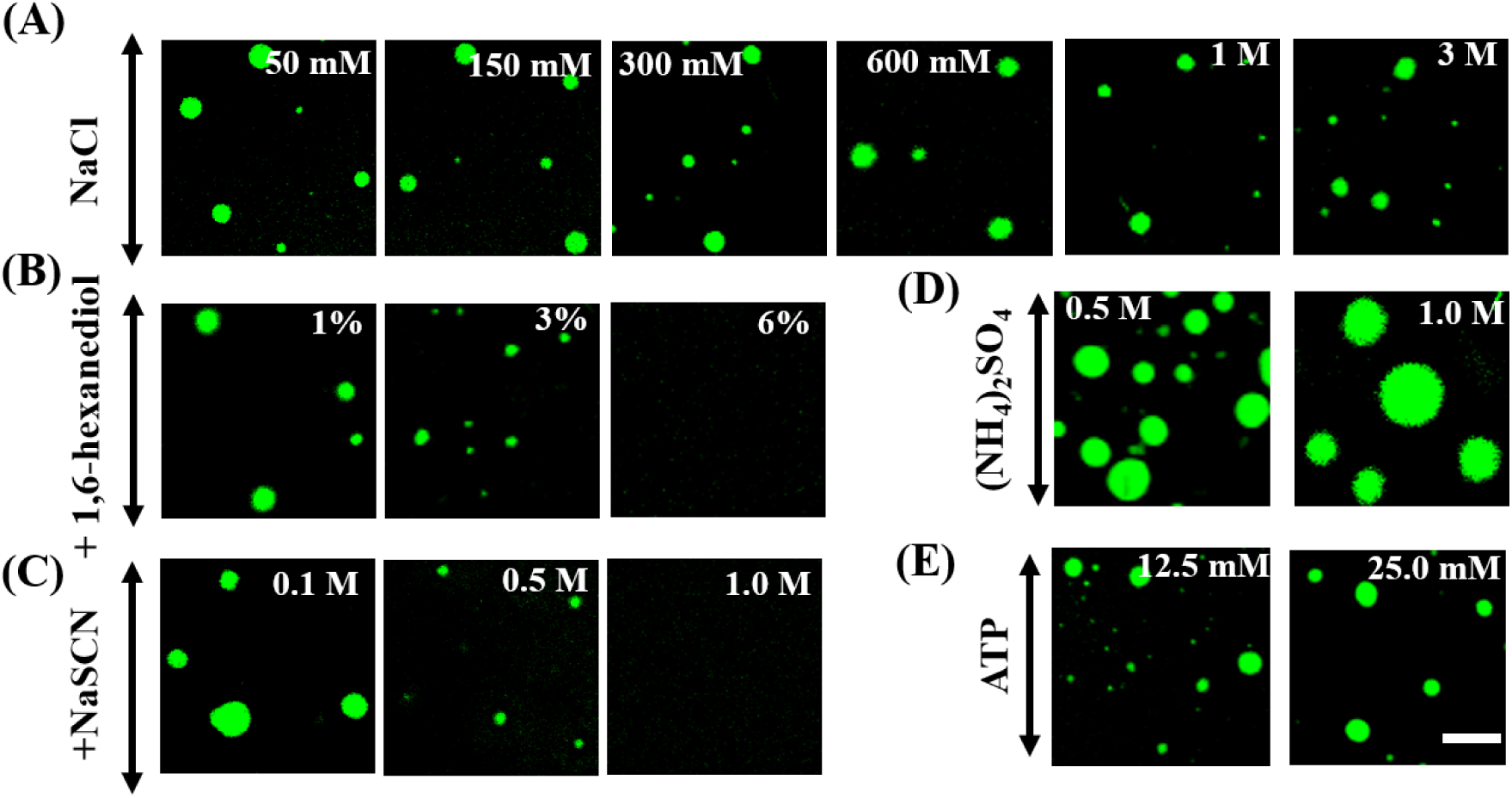
Confocal images showing the effect of concentrations of (A) NaCl, (B)1,6-hexanediol, (C) NaSCN, (D) (NH_4_)_2_SO_4_, and (E) ATP on the stability of FITC-labeled trypsin (5 nM) droplets at 37 °C in pH 7.4 PBS. Scale bar corresponds to 5 μm.

### Ca^2+^-Binding Reversibly Regulates the LLPS of Trypsin-Like Serine Proteases

Having established that the phase separation of trypsin is a spontaneous process in the presence of inert crowders through the involvement of multivalent hydrophobic interactions between short patches of polypeptide chains, we seek to know how this spontaneous process influence the catalytic activity of trypsin and trypsin-like serine proteases in the absence and presence of Ca^2+^. It is a known fact that Ca^2+^ binds at the CBL of trypsin and enhance its catalytic activity and stability.^22–27,34^ However, apart from these two functional roles, no other physiological function of Ca^2+^ is known for trypsin-like serine proteases. First, we seek to address whether Ca^2+^ binding influences the spontaneous LLPS of trypsin in the presence of inert crowders (Figure 5A). Notably, this fundamental question has not been addressed previously as the present study is first of its kind to discover the spontaneous LLPS of trypsin-like serine proteases under crowding environment. The phase separation assays were monitored under confocal microscope using different concentrations (1–10 mM) of Ca^2+^-equilibrated trypsin (5 µM) in the presence of 10% PEG at 37 ℃ in pH 7.7 tris-HCl buffer. CLSM images reveal remarkable influence of Ca^2+^ on the feasibility of LLPS of trypsin. While the presence of 1 mM Ca^2+^ significantly reduces the size of trypsin droplets, addition of 5 and 10 mM Ca^2+^ completely prevents the phase separation of trypsin (Figure 5B). Similar as Ca^2+^, 10 mM Mg^2+^ also prevents the LLPS of trypsin (Figure S6). To know whether this process is reversible or not, the trypsin bound Ca^2+^ ions (10 mM) were first removed upon addition of different concentrations of EDTA into the mixture and finally the phase separation was initiated upon addition of 10% PEG (Figure 5C). Interestingly, increasing the EDTA concentration from 1 to 10 mM results in the reappearance of trypsin droplets as revealed from the CLSM images (Figure 5D). It should be noted that the mean droplet size of trypsin increases with increasing the concentration of EDTA under the same experimental conditions due to gradual removal of trypsin bound Ca^2+^ and complete removal occurs at 10 mM EDTA. To know whether Ca^2+^ can able to bind with the phase-separated trypsin and similar reversibility establishes in the presence of EDTA, we formulated a phase separation assay as depicted in Figure 5E. Initially, trypsin was allowed to phase separate in the presence of 10% PEG at 37 ℃. CLSM image confirms the presence of well-dispersed spherical droplets of trypsin (Figure 5F(ii)). To this phase-separated droplets of trypsin, we sequentially add 10 mM Ca^2+^ followed by 10 mM EDTA. We envision that as trypsin retains its native-like conformation upon phase-separation, they will spontaneously bind with Ca^2+^ and show similar reversible transition in the presence of EDTA. Interestingly, addition of 10 mM Ca^2+^ into the phase-separated trypsin results in complete disappearance of droplets (Figure 5F(iii)), suggesting spontaneous binding of Ca^2+^ with the phase-separated trypsin. More importantly, subsequent addition of 10 mM EDTA results in the reappearance of phase-separated droplets of trypsin (Figure 5F(iv)). Notably, α-chymotrypsin also exhibits similar reversible modulation of LLPS in the presence of Ca^2+^ and EDTA (Figure S7). Therefore, our findings reveal that Ca^2+^ ions can effectively modulate the phase separation of trypsin-like serine proteases.

**Figure 5.**
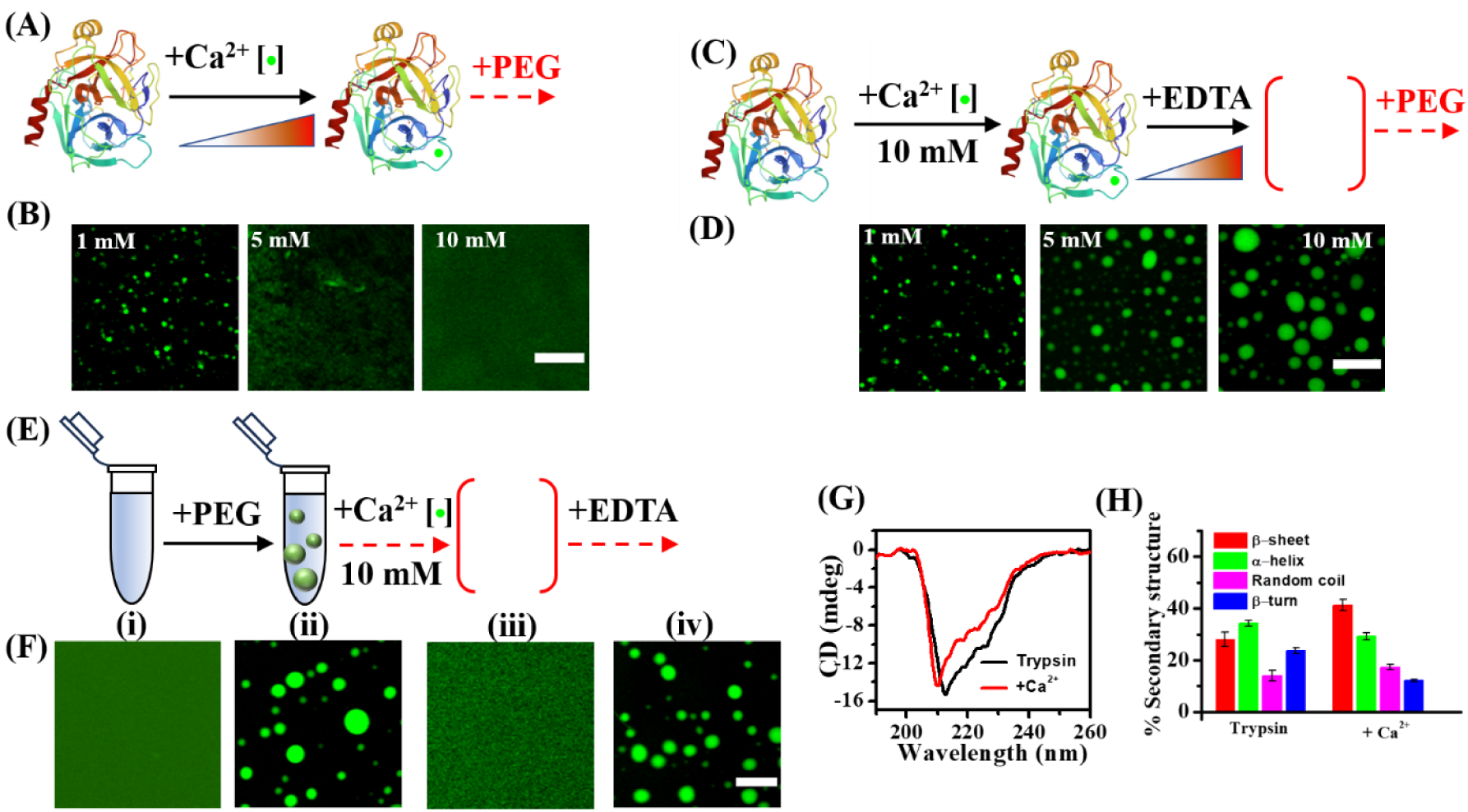
(A) Schematic and (B) confocal images showing the effect of Ca^2+^ concentrations on the feasibility of phase separation of 5 µM trypsin in the presence of 10% PEG 8000 at 37°C in pH 7.7 tris-HCl buffer. (C) Schematic and (D) confocal images showing the effect of EDTA concentrations on the feasibility of phase separation of Ca^2+^-bound trypsin in the presence of 10% PEG 8000. (E) Schematic and (F) confocal images showing the effect of 10 mM Ca^2+^ and 10 mM EDTA on the phase-separated droplets of FITC-labeled 5 µM trypsin. (G) CD spectra and (H) secondary structure contents of 100 µM trypsin in the absence and presence of 2 M Ca^2+^. The data points represent the mean ± s.e.m. from three independent measurements. Scale bars correspond to 5 µm.

The absence of any phase separation in the presence of Ca^2+^ suggests conformational alteration of native trypsin which ultimately prevents intermolecular protein-protein interactions. To validate this possibility, we recorded CD, FTIR, and Raman spectra of trypsin in the absence and presence of Ca^2+^. The CD spectrum of Ca^2+^ bound trypsin exhibits significant spectral changes compared to that of free trypsin (Figure 5G). The CD peak at 213 nm of free trypsin shifts to 210 nm upon binding with Ca^2+^. Moreover, the spectral intensity of Ca^2+^ bound trypsin decreases noticeably compared to that of free trypsin. The deconvoluted FTIR and Raman spectra reveal noticeable increase in the β-sheet and random coil content of Ca^2+^-bound trypsin at the expense of α-helix and β-turn (Figures 5H and S8). The β-sheet content increases from 28 to 42% and the α-helix and β-turn contents decrease from 34 to 29% and 24 to 12%, respectively upon Ca^2+^ binding. Notably, similar increase in the β-sheet content and decrease in the α-helix content have been observed previously for free trypsin at lower acidic pH of 3.5. Taken together, our present findings reveal that the inhibitory effect of Ca^2+^ on the LLPS of trypsin is primarily due to the conformation changes of Ca^2+^-bound trypsin which prevents the protein-protein interactions. Finally, we aimed to address how this regulatory role of Ca^2+^ influences the catalytic activity of free and phase-separated trypsin and whether this regulatory effect has any physiological relevance in the context of cellular function of trypsin-like serine proteases.

### The Role of Ca^2+^ and LLPS on the Esterase Activity of Trypsin

In the present study, we studied the trypsin-catalyzed hydrolysis of *p*-nitrophenyl acetate (*p*-NPA) to *p*-nitrophenolate (*p*-NP) as a model reaction (Figure 6A). The *p*-NPA to *p*-NP conversion can be conveniently followed using UV-vis spectroscopy by monitoring the spectral signature of *p-*NP at 400 nm.^34^ We performed all the kinetic experiments in pH 7.7 tris-HCl buffer at 37 ℃ using 7 µM trypsin.

**Figure 6.**
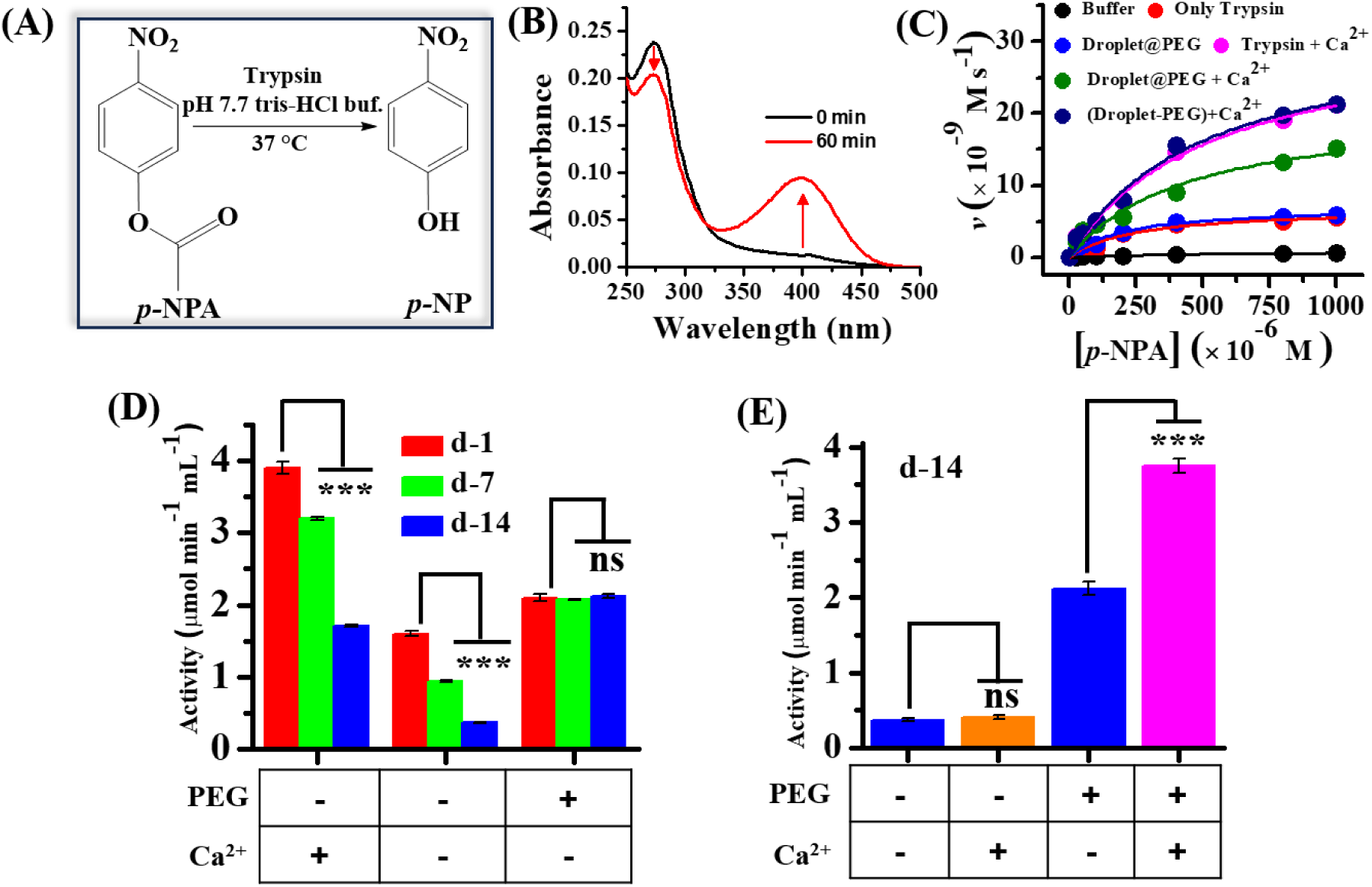
(A) Schematic showing the trypsin catalyzed conversion of *p*-NPA to *p*-NP at 37 °C in pH 7.7 tris-HCl buffer. (B) Changes in the UV-vis absorption spectra of the reaction mixture containing 25 µM *p*-NPA and 5 µM trypsin in pH 7.7 tris-HCl buffer upon 60 min of reaction. (C) Michaelis-Menten plots of 7 µM trypsin catalyzed hydrolysis of *p*-NPA under different conditions. (D) Effect of aging on the activity of 5 µM trypsin in the absence and presence of 10% PEG 8000 and 10 mM Ca^2+^ for a period of 14 days. (E) Effect of 10 mM Ca^2+^ on the activity of free trypsin and phase-separated trypsin samples upon 14 days of aging. The data represent the mean ± s.e.m. from three independent experiments. Statistical significance was assessed by a two-tailed, unpaired Student’s *t*-test with ***, *P* value < 0.001; and not significant (ns), P ˃ 0.05. All the measurements were performed in pH 7.7 tris-HCl buffer at 37 °C.

Addition of 7 µM trypsin into the 25 µM solution of *p*-NPA in pH 7.7 tris-HCl buffer at 37 ℃ results in the appearance of a distinct absorption peak at 400 nm after 60 min of reaction (Figure 6B). The esterase activities of trypsin at different conditions were monitored by recording the changes in the absorbance at 400 nm as a function of reaction time (Figure S9). The enzymatic rate of trypsin follows a typical Michaelis-Menten behavior as a function of *p*-NPA concentrations (Figure 6C). In the absence of trypsin, no noticeable conversion of *p*-NPA to *p-* NP has been observed, indicating the catalytic role of trypsin towards the hydrolysis of ester bond. Notably, the rate increases drastically in the presence of 13 mM Ca^2+^ relative to that of Ca^2+^-free trypsin. These kinetic data were fitted with the Michaelis-Menten equation to estimate the maximum velocity (*V*max) and Michaelis constant (*K*_m_). The estimated values of *V*_max_, *K*_m_, and turnover numbers (*k*_cat_) are tabulated in Table S1. While the *k*_cat_ of trypsin increases by a factor of 4.63 to 4.40 × 10^-3^ s^-1^ in the presence of Ca^2+^, the *K*_m_ increases by a factor of 2.1 to 5.28 × 10^-4^ M. These kinetic paraments of Ca^2+^-bound trypsin match well with those reported previously.^34^ The increase in the *K*_m_ value of trypsin in the presence of Ca^2+^indicates lower binding affinity towards *p*-NPA possibly due to the conformational alteration as revealed from our CD and FTIR measurements.

Next, we explore how the presence of PEG alters the esterase activity of trypsin in the absence and presence of Ca^2+^. As PEG promotes phase separation of trypsin, we first equilibrated trypsin with 10% PEG for 1 h. The reaction kinetics were monitored just after the addition of *p*-NPA into the phase-separated trypsin. Notably, the Michaelis-Menten plot and estimated kinetic paraments for phase-separated trypsin match well with those of free trypsin in the absence of Ca^2+^ (Figure 6C and Table S1), suggesting that phase-separated trypsin retained its native-like conformation and PEG has minimal effect on the kinetics of this reaction. However, the *k*_cat_ value estimated for Ca^2+^-bound trypsin in the presence of PEG shows a value of 3.01 × 10^-3^ s^-1^, which is 1.5 times lower than that estimated for Ca^2+^-bound trypsin in the absence of PEG (Table S1). As Ca^2+^ prevents phase separation of trypsin in the presence of PEG, the differences observed in the turnover number could be possibly due to the presence of PEG in the reaction mixture. It should be noted that polymeric crowders are known to retard the catalytic rates of various enzymes including trypsin.^34,36,38,43,71^ Moreover, it is well documented in the literature that polymeric crowders increase the viscosity of the reaction mixture which ultimately slowdown the rate of diffusion-controlled enzymatic reactions.^71^ To validate this possibility, we first allowed trypsin to undergo phase separation upon incubation with 10% PEG for 1 h. Subsequently, the dilute phase containing PEG was removed from the dense droplet phase of trypsin by simple centrifugation, followed by redispersion of the droplet phase in buffer. To this PEG free dispersion of trypsin droplets, 13 mM Ca^2+^ was added and the esterase activity was monitored after the addition of *p*-NPA. Interestingly, after the removal of PEG, the Michaelis-Menten plot exactly overlaps with that of Ca^2+^-bound trypsin alone in the absence of PEG (Figure 6C). Moreover, the estimated kinetic parameters match well with those estimated for Ca^2+^-bound trypsin alone (Table S1). Taken together, these findings indicate that both free and phase-separated trypsin can bind with Ca^2+^ and exhibits comparable esterase activity under physiological conditions.

Finally, we seek to know whether the observed biomolecular condensation of trypsin-like serine proteases has any physiological role in regulating their enzymatic activity. More specifically, we want to understand the specific role of this spontaneous and natural condensation phenomenon on the autolysis, premature activation, and Ca^2+^ signalling processes which are intrinsically associated with the enzymatic pathways of trypsin-like serine proteases. To address this question, we monitored the activity of trypsin over a period of 14 days in the absence and presence of PEG and Ca^2+^ using UV-vis spectroscopy (Figure S10). In the absence of PEG, the enzymatic activity of free and Ca^2+^-bound trypsin decreases significantly over a period of 14 days (Figure 6D). This decrease in activity as a function of aging can be explained by considering autolysis of both Ca^2+^-bound and -free trypsin. Notably, the autolysis rate is slower in the presence of Ca^2+^ as reported previously.^12,23–25^ Autolysis is a major deactivation/degradation pathway for trypsin-like serine proteases as described in previous studies.^12,23–25^ In contrast, we observed an interesting trend in the case of phase-separated trypsin in the presence of 10% PEG. It is observed that the phase-separated droplets of trypsin can effectively preserve its initial enzymatic activity over a period of 14 days without any noticeable loss. This novel behavior can only be explained if we consider negligible autolysis of the phase-separated trypsin. The unique conformational stabilization of trypsin inside the droplets prevents them to undergo autolysis. Furthermore, to know how and whether the 14 days aged samples of free trypsin and phase-separated trypsin respond to Ca^2+^-dependent activation, we compared their enzymatic activity after 14 days of aging (Figure 6E). While the free trypsin sample (d-14) shows no Ca^2+^-dependent enhancement of its activity, remarkable boosting (9-fold) in the enzymatic activity of phase-separated trypsin sample (d-14) has been observed in the presence of Ca^2+^. These observations clearly reveal that the phase-separated droplets of trypsin can preserve/protect the native-like active conformation and enzymatic activity for a prolong period and whenever required, they can be activated for catalytic transformations via Ca^2+^ signalling. Based on our findings, we present a schematic model where living cells synchronize the precise activation of trypsin-like proteases via several regulatory steps to avoid autolysis and premature trypsin activation which may leads to pancreatic self-digestion (Scheme 2). It is quite evident that cells utilize biomolecular condensation of trypsin-like serine proteases to store and preserve the native-like unique conformation to prevent autolysis and whenever required, they can be activated at the right time and at right place. To the best of our knowledge, the present study is the first report to showcase this unique and hidden role of biomolecular condensates of trypsin-like serine proteases and may finds wide-ranging implications in cell physiology and biotechnological applications.

**Scheme 2.**
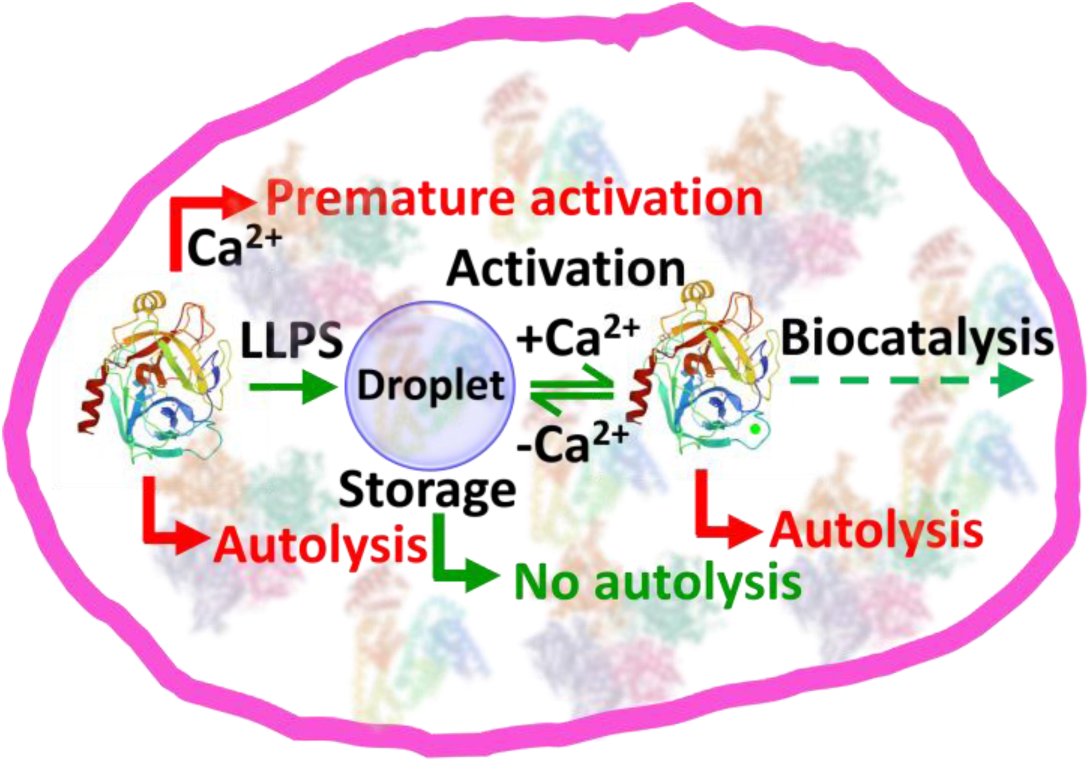
Schematic Illustration Showing the Spatio-Temporal Regulation of the Activity of Trypsin-Like Serine Proteases via Synchronization of the LLPS with Various Activation and Deactivation Pathways

### Summary

In the present study, we discovered a novel mechanistic pathway by which cells might regulate the enzymatic activity of trypsin-like serine proteases in a spatio-temporal manner. We have shown that trypsin-like serine proteases undergo spontaneous LLPS under macromolecular crowding via multivalent hydrophobic interactions in a broad range of protein concentrations, pH, temperature, and ionic strengths. The phase-separated droplets of trypsin retained its native-like active conformation and do not undergo liquid-to-solid-like phase transition. The phase separation of trypsin-like serine proteases has been shown to be reversibly regulated by Ca^2+^. Binding of Ca^2+^ prevents the phase separation of trypsin and α-chymotrypsin, whereas its removal with stoichiometric amount of EDTA reestablished the phase separation. These findings have been explained by considering significant conformation alteration of proteins upon binding with Ca^2+^ which prevent favorable protein-protein intermolecular interactions. More importantly, we have illustrated that the phase-separated droplets of trypsin can preserve its esterase activity over a period of 14 days without any autolysis, whereas free trypsin significantly loses its activity due to autolysis. Furthermore, the phase-separated droplets of trypsin spontaneously respond to Ca^2+^-mediated activation of esterase activity even after 14 days of aging, which is not feasible for free trypsin. The present study paves the way for further in-depth understanding of the hidden mechanistic pathways of other serine proteases to better understand their role in cell physiology and utilization toward industrial applications.

## ASSOCIATED CONTENT

### Supporting Information

Materials, experimental methods and characterization techniques; confocal image of FITC-labeled trypsin in the absence of PEG; confocal images of FITC-labeled porcine pancreatic trypsin and α-chymotrypsin in the presence of PEG; confocal image of FITC-labeled trypsin in the presence of 20 mg/mL BSA; confocal image and intensity line profiles of trypsin droplets with FITC-labeled 20 mg/mL BSA; FTIR spectra of trypsin in the absence and presence of different crowders; confocal image and FTIR spectra of trypsin in the presence of Mg^2+^; confocal images of FITC-labeled trypsin droplets followed by the addition of Ca^2+^and EDTA; Raman spectra and percentage secondary structure estimated from the deconvoluted Raman spectra of trypsin in the absence and presence of Ca^2+^; Plots of absorbance at 400 nm against the reaction time at different concentrations of *p*-NPA in different conditions; Changes in the UV-vis absorption spectra of 25 μM *p*-NPA in the presence of 5 µM trypsin at different reaction conditions as a function of aging; Table of kinetic parameters of trypsin-catalyzed esterase reaction under different conditions; and references. (PDF)

## Notes

The authors declare no competing financial interests.

## Acknowledgements

The authors acknowledge Indian Institute of Technology (IIT) Indore for providing financial support, and infrastructure. This work is financially supported by Council of Scientific and Industrial Research grant no. 01/3108/23/EMR-II. The authors acknowledge SIC, IIT Indore, for instrumental facilities. The authors thank Prof. Rajesh Kumar and Mr. Deb Kumar Rath for Raman measurements. The Raman facility received from the Department of Science & Technology (DST), Government of India, under the FIST scheme (Grant no. SR/FST/PSI-225/2016) is gratefully acknowledged. C. K. P. acknowledges Ministry of Education (MoE), India for research fellowships.

