## Supplemental file for "Biomolecular Condensation of Trypsin Prevents Autolysis and Promotes Ca^2+^-Mediated Activation of Esterase Activity"

### Table of Contents


| S.N. | | Page No. |
|---|---|---|
| 1 | Materials | S3 |
| 2 | Methods | S3-S6 |
| 3 | Characterizations | S6-S8 |
| 4 | Supporting Figures | S9-S18 |
| 5 | Table-1 | S19 |
| 6 | References | S20-S21 |

#### 1. Materials

Trypsin from bovine pancreas (essentially salt-free), polyethylene glycol 8000 (PEG 8000), mPEG-$NH_2$ (MW 5000), Ficoll 400, bovine serum albumin (BSA), dextran 70, sodium thiocyanate (NaSCN), 1,6-hexanediol, fluorescein-5-isothiocyanate (FITC), adenosine triphosphate (ATP), Hellmanex III and the Pur-A-Lyzert dialysis kit (molecular weight cutoff 3.5 kDa) were purchased from sigma-Aldrich. Porcine pancreatic trypsin, α-chymotrypsin from bovine pancreas, and *p*-nitrophenyl acetate were purchased from SRL. Sodium dihydrogen phosphate monohydrate ($NaH_2PO_4.H_2O$), di-sodium hydrogen phosphate heptahydrate ($Na_2HPO_4.7H_2O$), sodium acetate trihydrate ($CH_3COONa.3H_2O$), acetic acid ($CH_3COOH$), sodium bicarbonate ($NaHCO_3$), sodium carbonate anhydrous ($Na_2CO_3$), tris(hydroxymethyl)aminomethane (Tris buffer), sodium chloride (NaCl), ammonium sulphate ($(NH_4)_2SO_4$), magnesium sulphate ($MgSO_4$), hydrochloric acid (HCl), calcium chloride dihydrate ($CaCl_2.2H_2O$), ethylenediamine tetra acetic acid (EDTA) and methanol (MeOH) were purchased from Merck. All the chemicals were used without any further purification. Eco Testr pH1 pH meter was used to adjust the final pH (± 0.1) of all the buffer solutions. Milli-Q water was obtained from a Millipore water purifier system (Milli-Q integral).

#### 2. Methods

##### 2.1. Prediction of LCDs and IDRs of Trypsin

To predict the low complexity domains (LCDs) and disorder regions (IDRs) in trypsin, we used Simple Molecular Architecture Research Tool (SMART) (http://smart.embl-heidelberg.de/) and IUPred2 (https://iupred2a.elte.hu/), respectively.[1,2] The IUPed2 data were plotted using origin software.

##### 2.2. Preparation of Buffer Solutions and Crowder Solutions

Different buffer solutions with pH values of 3.5, 5, 7.4, 7.7, 9 and 10 were prepared by using Milli-Q water. The strength of each buffer was kept at 50 mM. Sodium citrate buffer (pH 3.5), sodium acetate buffer (pH 5), phosphate buffer saline (pH 7.4 PBS, 50 mM NaCl), tris buffer (pH 7.7, and 9), and carbonate-bicarbonate buffer (pH 10) solutions were used individually. All the buffer solutions were kept under sterile condition.

10% (w/v) PEG, 10% (w/v) dextran, 12.5% (w/v) Ficoll were prepared from a stock solution of 40% (w/v) PEG 8000, 40% (w/v) dextran 70, and 40% (w/v) Ficoll 400, respectively. 20 mg/mL BSA was prepared from the stock solution of 332 mg/mL BSA.

**2.3. Labeling of Enzymes and Crowders with Fluorescent Dyes**

The concentration of trypsin was estimated spectrophotometrically at $\lambda$ = 253 nm using the reported extinction coefficient of $8.80 \times 10^2$ $M^{-1}$ $cm^{-1}$,[3] and the concentration of α-chymotrypsin was measured using its absorbance at 280 nm with a molar extinction coefficient of $51 \times 10^3$ $M^{-1}$ $cm^{-1}$.[4] Trypsin and α-chymotrypsin were labeled with FITC dye according to an earlier reported method.[5] In short, 100 μM trypsin was mixed with FITC in a molar ratio of 1:10 ([trypsin]:[FITC]). The mixture was incubated for 4 h at room temperature followed by 6 h at 4 °C on a magnetic stirrer with slow rotation. After the completion of the reaction, the unconjugated dye molecules were removed using dialysis (molecular weight cut-off 3.5 kDa) against 50 mM PBS at 4 °C for 12 h with regular buffer exchange in 2 h intervals. The same procedure was followed for the labeling of mPEG-$NH_2$ (MW 5000) and BSA.

**2.4. LLPS Assay of Trypsin in the Presence of Crowders**

LLPS at different pH was examined by equilibrating 5 nM trypsin with 10% (w/v) PEG 8000 in different buffer solutions an equilibrated for different time intervals. The samples were prepared in 5 mL glass vials and were kept at 37 °C in the incubation chamber.

To check the time-dependent LLPS behaviour of trypsin, different concentrations of trypsin (0.005-5 μM) were prepared from a stock solution of 100 μM trypsin. The time-dependent droplet formation was confirmed by CLSM.

**2.5. Effect of Salts**

The salt-dependent phase separation assay was performed by adding different concentrations of NaCl (0-3 M), NaSCN (0-2 M), 1,6-hexanediol (0-10%), $(NH_4)_2SO_4$ (0.5 and 1 M), and ATP (12.5 and 25 mM) from their respective stock solutions. The solutions were prepared by adding 5 nM trypsin, 10% PEG 8000 and different concentrations of salts/aliphatic alcohol/ATP in pH 7.4 phosphate buffer. Next, the solutions were kept at 37 °C for 1 h and subsequently CLSM images were captured.

**2.6. Effect of Temperature and pH**

For the temperature-dependent study (4, 25, 70, and 90 °C), all the samples (5 nM trypsin in the presence of 10% PEG 8000) were prepared in pH 7.4 PBS and kept them for 1 h at different temperatures. Similarly, pH-dependent study was carried out using 5 nM trypsin in the presence of 10% PEG 8000 in different buffers (pH 3.5, 5, 7.7, 9, and 10) and kept for 1 h at 37 °C in incubation chamber.

**2.7. Effect of $Ca^{2+}$, $Mg^{2+}$, and EDTA**

To check the regulatory role of $Ca^{2+}$and $Mg^{2+}$, we prepared 5 μM trypsin in the presence of 1–10 mM $CaCl_2$ and 1–10 mM $MgSO_4$ from their respective stock solutions in pH 7.7 tris-HCl buffer at 37 °C. The stock solutions were prepared in pH 7.7 tris-HCl buffer. Then, we added 10% PEG 8000 into the solution and recorded CLSM images. To know the effect of EDTA, we added 1–10 mM EDTA solution into the reaction mixtures and recorded CLSM images.

**2.8. Enzyme Activity Study**

To examine the enzymatic activity of trypsin for a period of 14 days, we prepared samples of 5 μM trypsin in the absence and presence of $Ca^{2+}$ and 10% PEG and kept them inside a 37 °C incubation chamber. We performed trypsin-catalyzed conversion of 25 μM *p*-nitrophenyl acetate (*p*-NPA) to *p*-nitrophenol (*p*-NP) in pH 7.7 tris-HCl buffer at 37 °C with different samples and monitored trypsin activity at d-1, d-7, and d-14 using UV-vis spectroscopy. During the reaction, the absorbance of *p*-NP at $\lambda_{max}$ = 400 nm was monitored for a period of 60 min. The enzyme activity was estimated by using the following equation[6]

$$\text{enzyme activity } (\mu\text{mol min}^{-1}\text{mL}^{-1}) = \frac{(\Delta A)(\text{volume of reaction}) \times 100}{\varepsilon \times t \times V}$$

Here, $\Delta A$ is the difference in absorbance of test and blank sample at 400 nm, $\varepsilon$ is the molar absorption coefficient of *p*-NP at 400 nm ($\varepsilon$ = 18,000 $M^{-1}$ $cm^{-1}$),[7,8] $t$ is the total time taken for completion of the reaction, and $V$ is the volume of enzyme taken.

**2.9. Estimation of Kinetic Parameters**

The absorbance values obtained from UV–vis spectrometer was plotted as a function of time using OriginPro 8.1 software, and then the data were linearly fitted for each concentration. The initial rates were calculated by considering the molar extinction coefficients of *p*-NP ($\varepsilon_{400\ nm}$ = 18,000 $M^{-1}$ $cm^{-1}$). The estimated initial rates were plotted against substrate concentrations (25-1000 μM). The concentration of trypsin taken was 7 μM. For the removal of PEG from the phase separated droplets, the sample solution (7 μM trypsin + 10% PEG) was centrifuged at 18000 rpm for 20 min using Remi PR-24 centrifuge. The data were finally fitted with the Michaelis–Menten equation according to the following expression,

$$v = \frac{V_{max}[S]}{K_m + [S]}$$

where $v$ is the initial velocity, $[S]$ is the molar concentration of substrate, $V_{max}$ is the maximum velocity, and $K_m$ is the Michaelis constant. We used nonlinear curve fit analysis in OriginPro 8.1 software to estimate the fitted parameters $V_{max}$ and $K_m$. Finally, $k_{cat}$ was calculated by dividing the $V_{max}$ with the total enzyme concentration used.

**3. Characterization Techniques**

##### 3.1. UV–vis Spectroscopy

Absorption spectra and reaction kinetics were monitored by using PerkinElmer UV/vis/NIR spectrometer in a quartz cuvette (1 cm × 1 cm).

##### 3.2. Confocal Laser Scanning Microscopy (CLSM)

The confocal images were captured using an inverted confocal microscope, Olympus fluoView (model FV1200MPE, IX-83), through an oil immersion objective (100× 1.4 NA). The samples were excited using a diode laser (488 nm) by using appropriate dichroic and emission filters in the optical path. Samples for confocal measurements were prepared in liquid chambers. Typically, a 10 μL aliquot of the sample solution was placed onto a cleaned glass slide and sandwiched with a Blue Star coverslip. Finally, the sides of the coverslips were sealed with a minimal amount of commercially available nail paint, and then images were captured. The prepared samples were placed in a desiccator under vacuum.

##### 3.3. Fourier-Transform Infrared (FTIR) Spectroscopy

FTIR spectroscopy was performed to determine the secondary structure of protein using a Bruker spectrometer (Tensor-27). Liquid samples of 10 μL aliquots containing 100 μM trypsin in the absence and presence of different crowders, metal ions, and different pH solutions were used for FTIR measurements. The spectra were recorded in the range of 4000–400 $cm^{-1}$. Fourier self-deconvolution (FSD) method was used to deconvolute the spectra corresponding to the wavenumbers 1700−1600 $cm^{-1}$.[9] The Lorentzian curve fitting was performed to fit the spectra using origin 8.1 software. All the experiments were performed thrice with similar observations.

##### 3.4. Circular Dichroism (CD) Spectroscopy

CD spectra were recorded on a JASCO J-815 CD spectropolarimeter using a quartz cell of 1 mm path length with a scan range of 190-260 nm. Scans were recorded with a slit width of 1 mm and speed of 50 nm/min. The spectra were plotted using origin 8.1 software.

##### 3.5. Raman Spectroscopy

Raman spectra were recorded using Horiba-Jobin Yvon micro-Raman spectrometer with a 532 nm excitation laser and CCD detector. Protein samples (100 μM trypsin in the absence and presence of 2 M $Ca^{2+}$) of 15-20 μL aliquots were drop-casted on cleaned coverslips. A 532 nm excitation laser light with 40 mW (100%) laser power was utilized to excite the samples. The laser was focused using a 50× objective lens. The spot size of laser beam in the Raman spectrometer is ~2 μm and the diffraction grating of 600 lines/mm were used to disperse the scattering light. Raman data were collected in the backscattering mode with flat correction using inbuilt method in Raman spectrometer software. The obtained experimental data were deconvoluted in the amide-I region corresponding to the wavenumbers 1700−1600 $cm^{-1}$.

### Supporting Information Figures

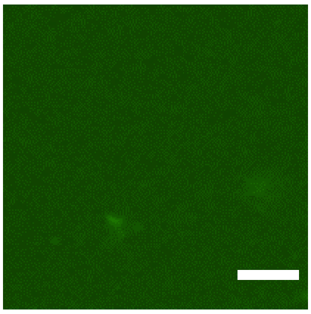

**Figure S1.** Confocal image of FITC-labeled 5 nM trypsin shows no droplet formation in the absence of PEG 8000 upon 1 h of incubation at 37 °C in pH 7.4 PBS. The scale bar corresponds to 5 μm.

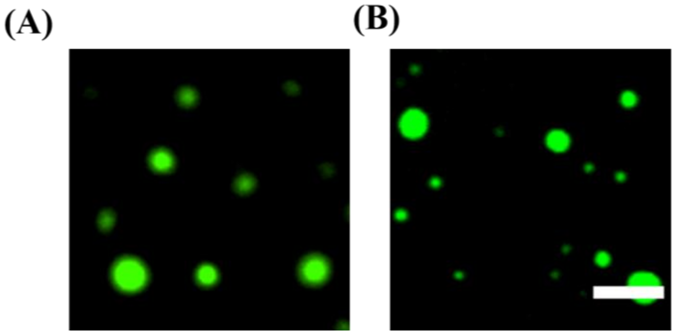


**Figure S2.** Confocal images of FITC-labeled (A) 5 nM porcine pancreatic trypsin, and (B) 5 nM α-chymotrypsin shows droplet formation in the presence of 10% PEG 8000 upon 1 h incubation at 37 °C in pH 7.4 PBS. The scale bar corresponds to 5 µm.

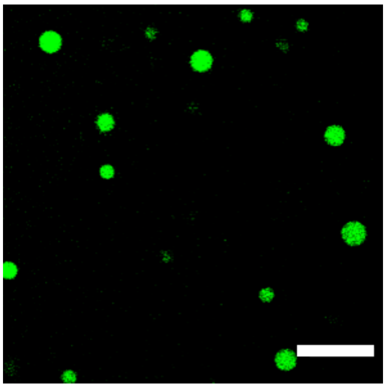

**Figure S3.** Confocal image of 5 nM FITC-labeled trypsin shows droplet formation in the presence of 20 mg/mL BSA upon 1 h incubation at 37 °C in pH 7.4 PBS. The scale bar corresponds to 5 μm.

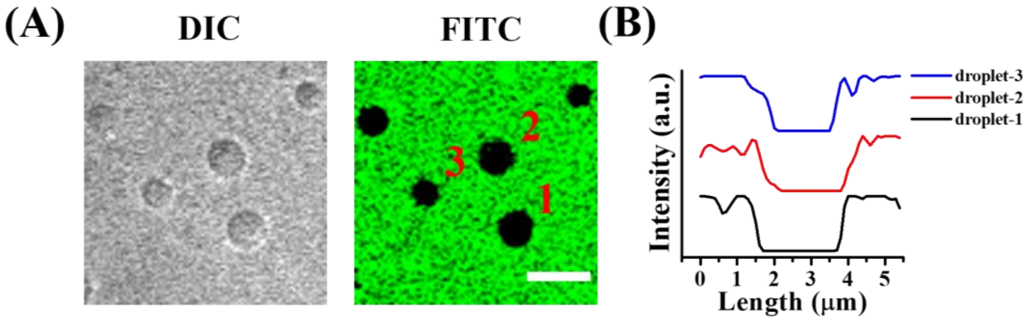


**Figure S4.** (A) Confocal images of unlabeled trypsin in the presence of 20 mg/mL FITC-labeled BSA, and (B) Intensity line profiles of three representative droplets. The scale bar corresponds to 5 μm.

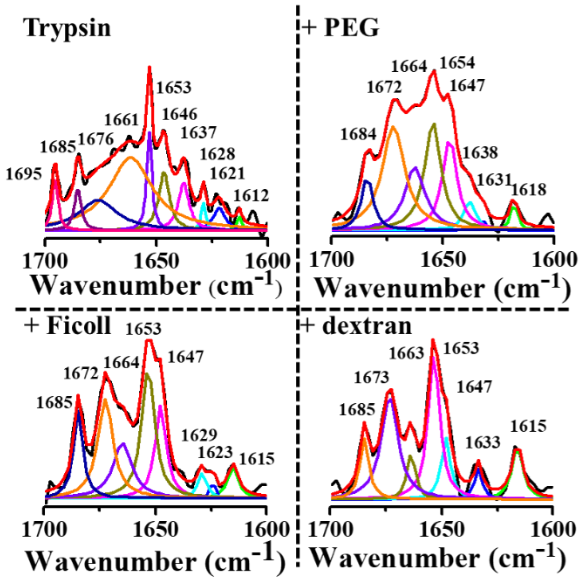


**Figure S5.** Deconvoluted FTIR spectra of 100 μM trypsin in the absence and presence of different crowders incubated for 1 h at 37 °C in pH 7.4 PBS. The experiments were performed three times with similar observations.

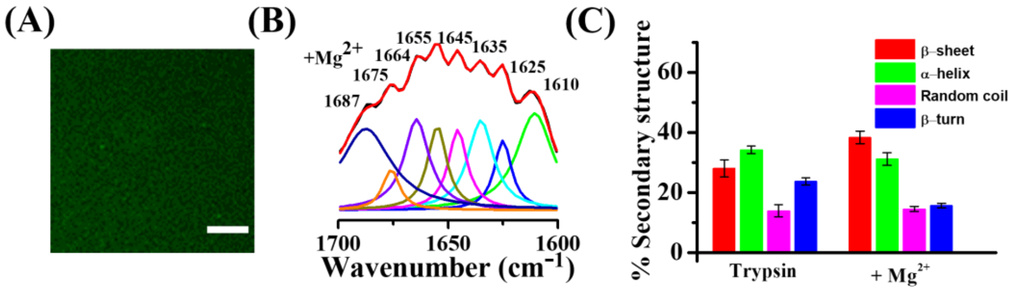


**Figure S6.** (A) Confocal image of FITC-labeled 5 μM trypsin showing no droplet formation with 10% PEG 8000 in the presence of 10 mM $MgSO_4$ in pH 7.7 tris-HCl buffer at 37 °C. (B) Deconvoluted FTIR spectrum of 100 μM trypsin in the presence of $MgSO_4$. (C) Secondary structure contents of 100 μM trypsin in the absence and presence of 2 M $Mg^{2+}$. The scale bar corresponds to 5 μm.

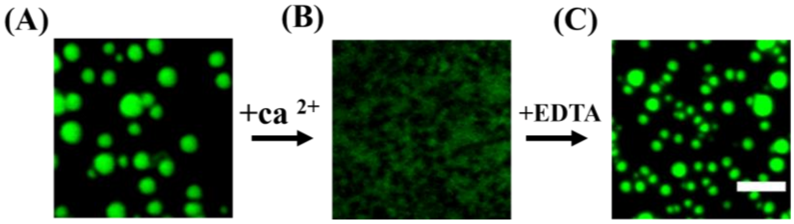


**Figure S7.** Confocal images of FITC-labelled 5 μM α-chymotrypsin recorded in the presence of (A) 10% PEG 8000, (B) 10% PEG 8000 and 10 mM $Ca^{2+}$, and (C) 10% PEG 8000, 10 mM $Ca^{2+}$, and 10 mM EDTA in pH 7.7 tris-HCl buffer at 37 °C. The scale bar corresponds to 5 μm.

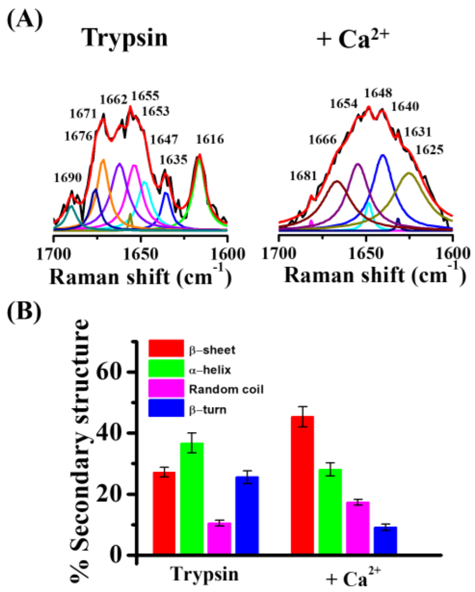


**Figure S8.** (A) Deconvoluted Raman spectra of 100 μM trypsin in the amide-I region in the absence and presence of 2 M $Ca^{2+}$ incubated at 37 °C for 1 h in pH 7.7 tris-HCl buffer. (B) Secondary structure contents of trypsin in the absence and presence of 2 M $Ca^{2+}$ calculated from the deconvoluted Raman spectra. The data points represent the mean ± s.e.m. for three independent measurements.

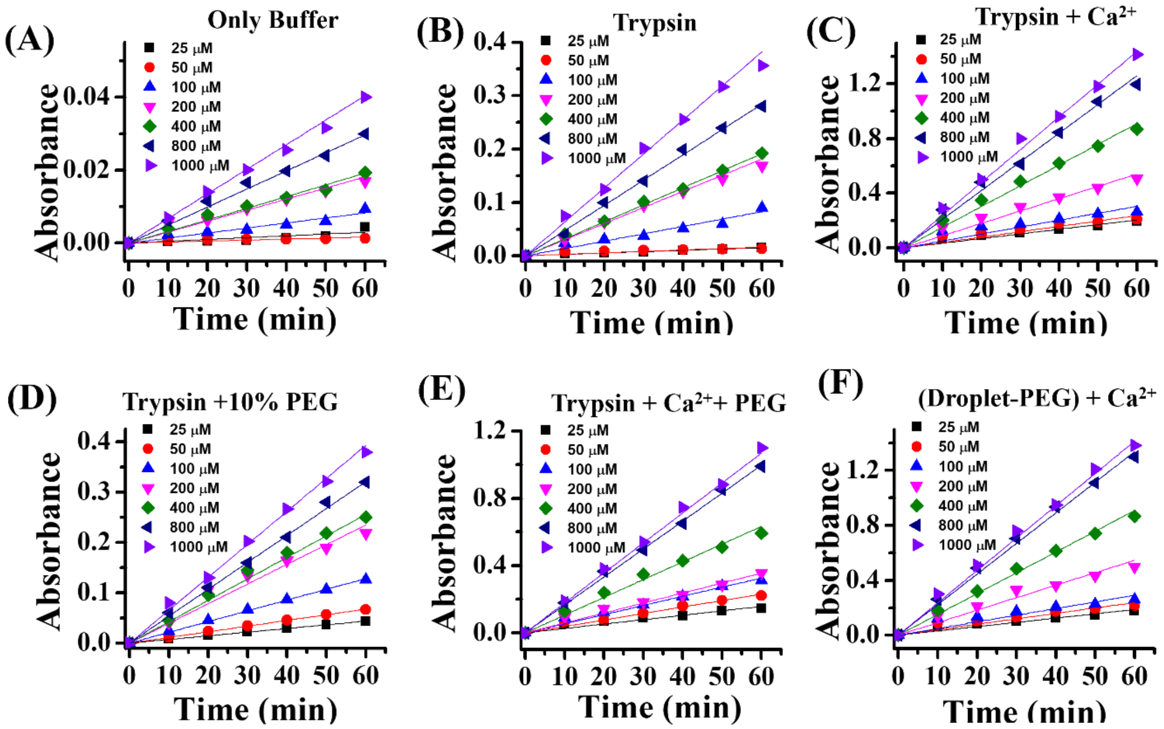


**Figure S9.** Plots of absorbance at 400 nm against the reaction time at different concentrations of *p*-NPA in (A) Buffer, (B) 7 µM Trypsin, (C) 7 µM Trypsin in the presence of 13 mM $Ca^{2+}$, (D) 7 µM Trypsin in the presence of 10% PEG, (E) 7 µM Trypsin in the presence of 13 mM $Ca^{2+}$ and 10% PEG and (F) PEG-separated droplets of trypsin in the presence of 13 mM $Ca^{2+}$ in pH 7.7 tris-HCl buffer at 37 °C.

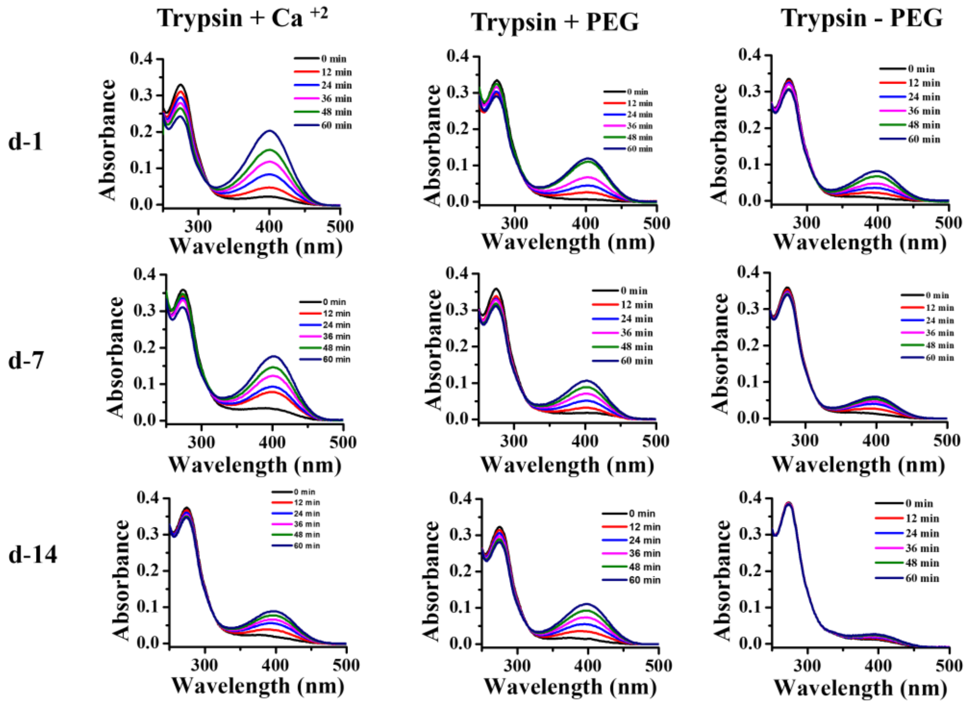


**Figure S10.** Changes in the UV-vis absorption spectra of 25 μM *p*-NPA in the presence of 5 μM trypsin at different reaction conditions as a function of aging (d-1, d-7, and d-14) in pH 7.7 tris-HCl buffer at 37 °C.

**Table S1. Kinetic Parameters of Trypsin-Catalysed Esterase Reaction Under Different Conditions**

| | $K_m$ (M) | $V_{max}$ (M $s^{-1}$) | $k_{cat}$ ($s^{-1}$) |
|---|---|---|---|
| Trypsin | $2.48 \times 10^{-4}$ | $6.85 \times 10^{-9}$ | $0.97 \times 10^{-3}$ |
| Trypsin + $Ca^{2+}$ | $5.36 \times 10^{-4}$ | $3.24 \times 10^{-8}$ | $4.63 \times 10^{-3}$ |
| Trypsin + PEG (Droplet@PEG) | $2.03 \times 10^{-4}$ | $7.09 \times 10^{-9}$ | $1.01 \times 10^{-3}$ |
| Trypsin + $Ca^{2+}$ + PEG | $4.49 \times 10^{-4}$ | $2.10 \times 10^{-8}$ | $3.01 \times 10^{-3}$ |
| (Droplet-PEG) + $Ca^{2+}$ | $4.96 \times 10^{-4}$ | $3.22 \times 10^{-8}$ | $4.60 \times 10^{-3}$ |

(8) Zhang, Z. Y.; Clemens, J. C.; Schubert, H. L.; Stuckey, J. A.; Fischer, M. W.; Hume, D. M.; Saper, M. A.; Dixon, J. E. Expression, Purification, and Physicochemical

Characterization of a Recombinant Yersinia Protein Tyrosine Phosphatase. *J. Biol. Chem.* **1992**, *267*, 23759–23766.

(9) Yang, H.; Yang, S.; Kong, J.; Dong, A.; Yu, S. Obtaining Information about Protein Secondary Structures in Aqueous Solution Using Fourier Transform IR Spectroscopy. *Nat. Protoc.* **2015**, *10*, 382–396.
